# From kin to spatial proximity conspecifics: social network reorganisation in developing juvenile vervet monkeys

**DOI:** 10.64898/2025.12.24.696368

**Authors:** Mawa Dafreville, Charlotte Canteloup, Fannie Beurrier, Erica van de Waal, Ebi Antony George

**Affiliations:** Department of Ecology and Evolution, University of Lausanne, Switzerland; Inkawu Vervet Project, Mawana Game Reserve, Swart Mfolozi, KwaZulu Natal, South Africa; Laboratory of Cognitive and Adaptive Neurosciences, CNRS, UMR 7364, University of Strasbourg, France; The Sense Innovation and Research Center, Lausanne, Switzerland; Centre for Functional Biodiversity, School of Life Sciences, University of KwaZulu-Natal, Pietermaritzburg, South Africa

**Keywords:** social development, social interactions, primates, social networks, multiplex networks

## Abstract

Navigating social relationships is a primary determinant of fitness in group-living animals and thus the transition of individuals from infancy to fully fledged group members is marked by concomitant changes in interactions. This developmental trajectory is particularly relevant in primates, where individuals must navigate a social landscape composed of different types of relationships and interactions while gaining independence from their mothers. Yet, little is currently known about how different layers of sociality, like proximity and grooming interactions, co-develop during the sensitive juvenile phase. Here, we harness multilayer network analysis on observations from 18 wild vervet juveniles (4-36 months of age) from two social groups to dissect the maturation of the juvenile social network. We quantify developmental trajectories across three complementary scales: maternal dyadic interactions, individual network centrality, and global network topology. We find that social maturation is defined not by simple network expansion, but by structural reorganisation. Specifically, juveniles shed their early bias toward mothers and kin, replacing these primary bonds by social interactions aligning with spatial proximity. This restructuring is further modulated by socio-demographic traits. Higher maternal rank confers access to influential partners, while females are more central than males in grooming networks, foreshadowing adult social roles. By charting the reorganisation of multiple social layers, this study unravels the processes underpinning the juvenile transition to independence in primates.

## Introduction

In group living animals, the ability to navigate social relationships is a fundamental driver of individual fitness, predicting resilience to stress, survival and reproductive success [1–3]. Across taxa, the juvenile period is a critical stage as individuals acquire the socio-cognitive skills necessary for establishing and maintaining social relationships that will ultimately influence their fitness-related outcomes (e.g., ravens: [4]; Siberian jays: [5]; meerkats: [6]; and a recent review: [7]). In primates, early social life is dominated by mother–offspring interactions [8–11], which gradually give way to increased involvement with peers and other group members [12,13]. This transition is not only shaped by age but also by the social environment into which juveniles are born. In matrilineal species such as vervet monkeys, maternal rank strongly dictates offspring’s social opportunities, with high-ranking mothers providing greater access to grooming partners and coalitionary support [14–17]. Consequently, juvenile social ontogeny can be viewed as a process with dual trajectories, one of gaining independence from mothers and the second of increasing involvement into a broader, pre-existing social structure.

The social inheritance hypothesis [18] provides a mechanistic framework for network acquisition, positing that juveniles passively replicate their mothers’ associations while building their network. However, empirical evidence challenges this deterministic view. In wild vervet monkeys, juveniles fail to mirror their mothers’ social networks, largely due to the temporal instability of those maternal ties [17]. Similar constraints can be seen in elephants, where network stability limits the direct transfer of associations (Goldenberg et al., 2016). These divergences from the inheritance model suggest that network ontogeny is driven by social niche construction, a process where individuals actively engineer their relationships to ensure stability [19,20]. Vilette and colleagues [21], for instance, captured this process in juvenile vervet monkeys, demonstrating that they build their own social networks through continuous adjustments. Over time, they consolidate a few strong ties while maintaining many weaker ones, forming a stable social “core” within a larger, flexible network. Collectively, these findings indicate that juveniles’ relational environments extend beyond the social patterns inherited from their maternal ties.

Primate sociality is inherently multidimensional, encompassing proximity, grooming, aggression and play networks [22–24]. This compels juveniles to navigate a complex social landscape of distinct, yet interconnected, behavioural layers. Empirical evidence confirms that these layers of social life mature at different rates. In chimpanzees, juveniles gradually shift from broad play networks to more selective grooming relationships, while proximity patterns remain relatively stable [25]. Grooming networks exhibit non-linear developmental trajectories that vary according to sex and grooming in bonobos, revealing that social development is behaviour-specific rather than uniform [26]. However, current research analyses these behavioural trajectories separately, ignoring the structural interdependence between them. The association between different layers imposes constraints on juvenile social development, particularly in matrilineal societies with stable dominance hierarchies, such as vervet monkeys. In these societies, dominance rank and kinship structure patterns of association and access to partners, with juveniles’ social opportunities emerging within these broader rank-based and kinship-based dynamics [24,27,28]. Contrasting with these inherited constraints is the developmental imperative for social learning: juveniles first rely on their mother as a model, then increasingly learn from the most efficient individuals, and later even adopt behaviours introduced by new group members [29]. Gaining a holistic perspective on juvenile social development, therefore, requires a framework capable of integrating and analysing kinship, proximity and social interactions-based networks simultaneously.

Multilayer social network analysis provides this framework, modelling distinct interactions as interconnected layers within a single unified network [30–33]. Multiplex networks are a type of multilayer networks where inter-layer connections are restricted to the same individual across each social ‘layer’ (e.g., proximity, grooming). This structure is ideal for tracking how an individual’s social role and strategy change across different relational contexts. While this powerful analytical tool is applied extensively in fields like epidemiology and neuroscience [34,35], its use in animal behaviour is nascent but has proven highly effective. It has been used to identify the cryptic reproductive heir in social wasps [36], determine which females are central across multiple interaction types in rhesus macaques [37] and uncover how changes in a male’s position within grooming networks influence the aggression he receives in vervet monkeys [38]. However, this approach has been rarely leveraged to capture temporal changes in network structure, such as the dynamic restructuring that defines juvenile social development.

In this study, we dissect vervet juvenile social development across three fundamental scales of organisation. First, at the dyadic level, we investigate the process of maternal independence. We quantify this process by examining how maternal interactions change with age, predicting that the mother will gradually cease to be the primary interaction partner as juveniles mature [25,26]. Next, we scale up to the nodal (individual) level in juvenile egocentric networks to assess how this growing autonomy impacts a juvenile’s overall social position [8]. Using multilayer centrality metrics, we test how a juvenile’s role changes with age and how it is influenced by socio-demographic factors (i.e., sex, maternal rank, social group). We predict that centrality will increase with maternal rank, because juveniles of high-ranking mothers’ benefit from greater social opportunities (reflecting inherited social capital) and be higher for females, who remain in their natal group (reflecting female philopatry; [25,39]). Finally, at the global network level, we examine the structural architecture emerging from these changes. Using edge overlap analysis in multilayer networks, we test how the alignment between kinship, proximity, and social interaction networks changes with age. We hypothesise a social shift, predicting that the overlap of kinship networks with social interaction networks will decrease as juveniles gain independence, while the overlap between spatial proximity and social interaction networks increases. By integrating these three scales within a multilayer framework, our study provides the first unified model of primate social ontogeny.

## Method

### Population

The vervet monkeys aged from 4 to 36 months, were sampled from a free-range population of vervet monkeys (*Chlorocebus pygerythrus*) at the INKAWU Vervet Project (IVP) in Mawana Game Reserve in South Africa. The study has been conducted on 18 juveniles from two habituated groups: Kubu (KB, with 26 group members) and Ankhase (AK, with 22 group members; Table 1). Individuals were classified into three age classes: (i) less than 1 year old ([<1yo], weaning period, Fairbanks & Mcguire, 1995); (ii) 1 to 2 years old ([1yo], putative period of competition with a new younger sibling and (iii) 2 to 3 years old ([2yo], pre-independence stage), observed respectively from 4 to 11, 16 to 23 and 28 to 35 months of age [40–42]. Kinship was determined using matrilineal descent, with individuals belonging to one of the two main matrilines within each group.

**Table 1.**
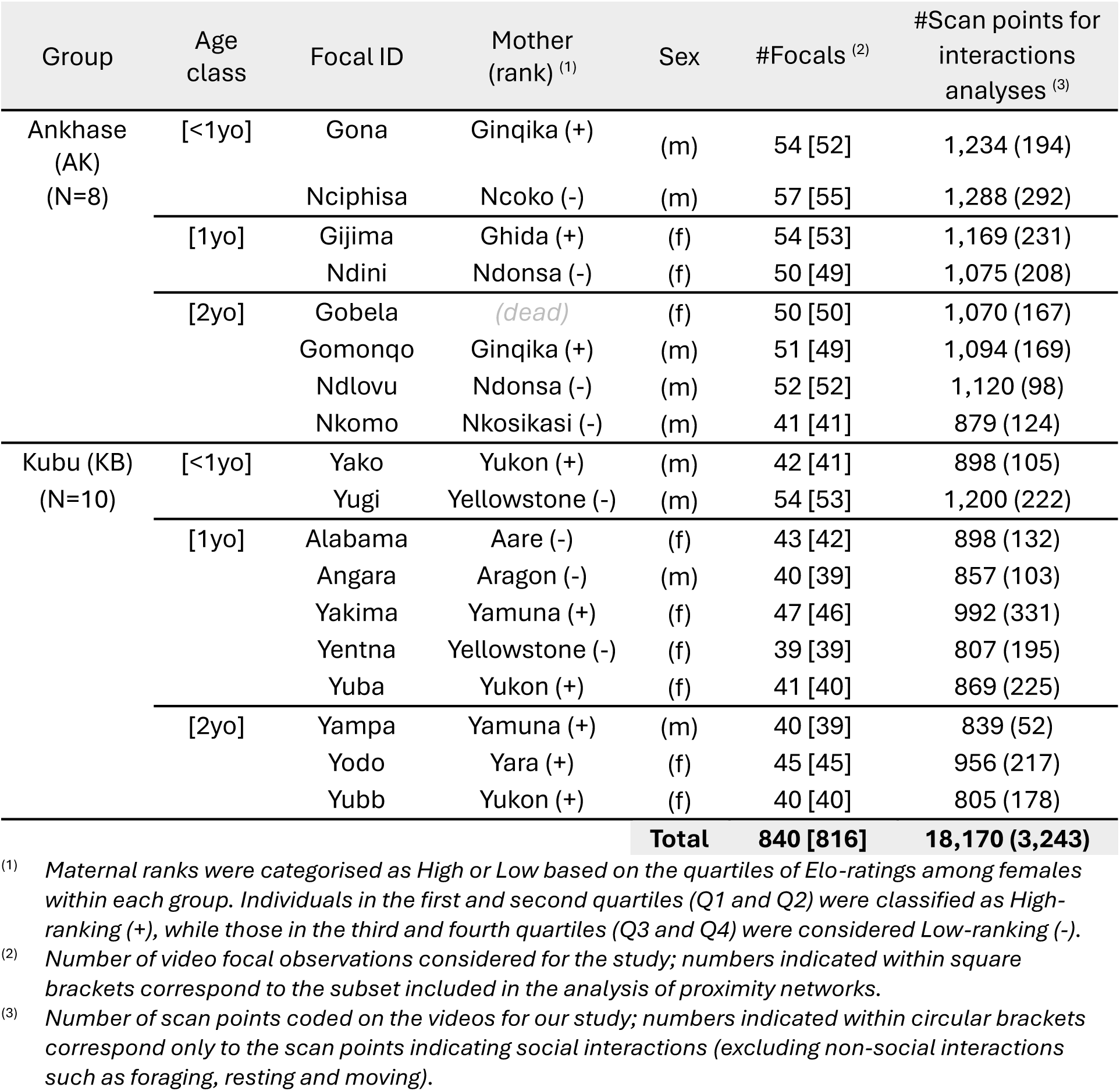
Number of focal observations and scan points per individual, by group and age category.

### Data collection

We filmed juveniles in their natural setting from March to October 2024 across three seasons: late-summer (March-April), early winter (July-August) and late-winter (September-October). All observations were conducted using a Panasonic HC-V770 camera equipped with a Rode VideoMic Pro Rycote Micro external microphone to record behaviour. Focal individuals were approached on foot to approximately 5 metres to record their behaviour in 10-minute sessions. To avoid repeated sampling, we followed a randomised order of individuals and tried to ensure that no focal individual was observed more than once per day and time-period (morning, mid-day, afternoon). To maximise the observations of social interactions, recordings were only initiated when the focal juvenile was within 5 metres of at least one other conspecific. A total of 840 ten-minute focal sessions were recorded (Table 1; AK: 40.95 ± 7.59 per individual, KB: 43.1 ± 4.58 per individual).

### Behavioural sampling and coding

We used a scan sampling method for obtaining two different types of data: (i) a proximity scan conducted at the beginning of each focal observation to orally record the identities of individuals within 1 meter radius of the focal animal; and (ii) a scan sampling method (Altmann, 1974) to record the activities of infants and juveniles every 30 seconds, along with additional relevant information (see behavioural repertoire in Table 2), using the coding software ELAN (Version 6.4). For each scan point assigned to the focal individual, we noted (1) the broad activity type, (2) when the activity was social and its type, (3) the presence of physical contact and (4) the identity and demographic details of any interaction partner.

**Table 2.**
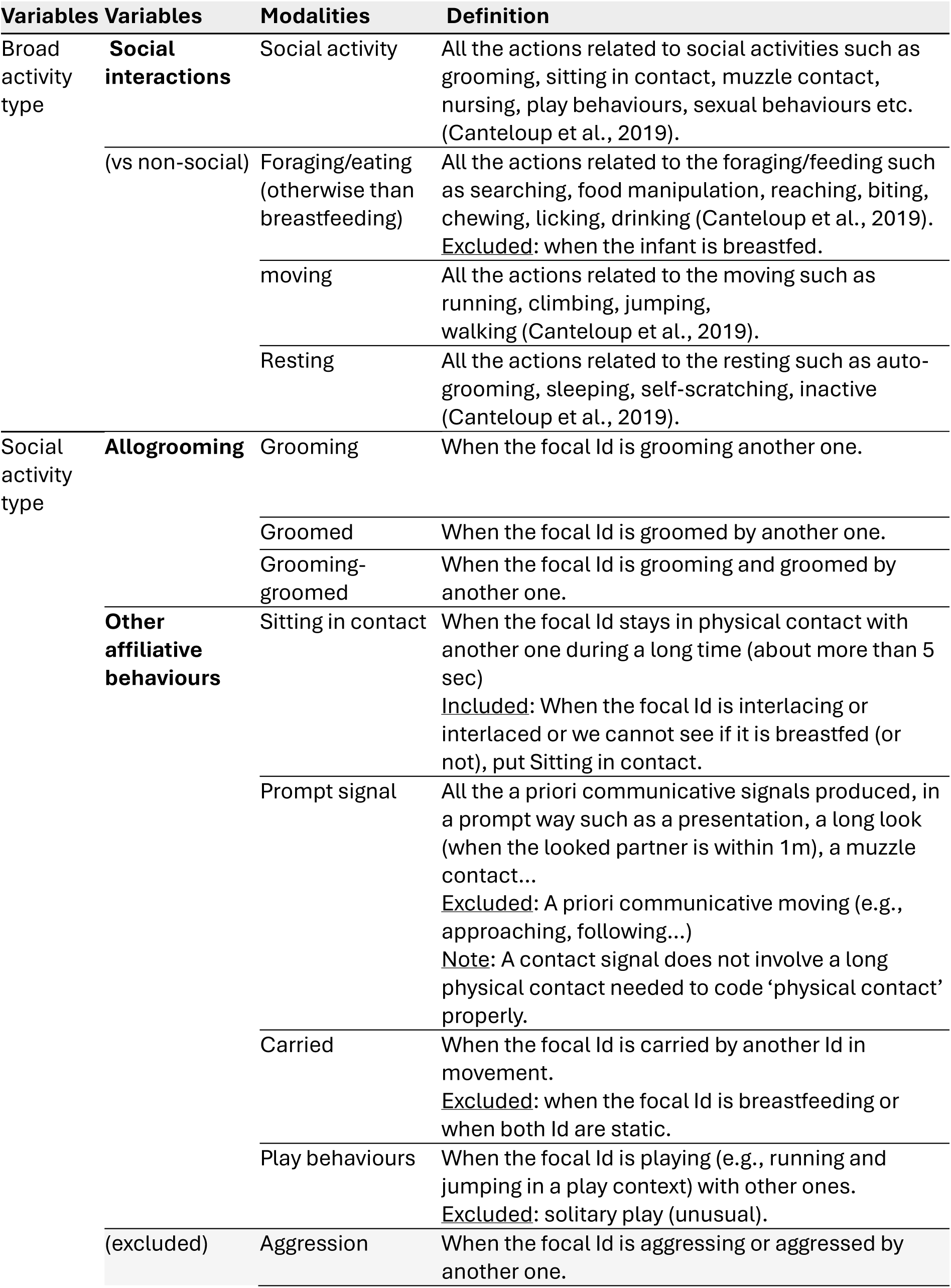
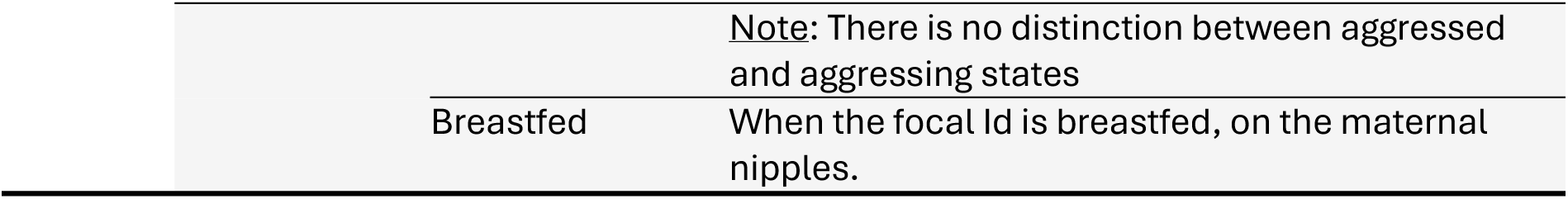
Behavioural repertoire of social interaction coding in juvenile vervet monkeys.

After coding the video dataset (based on Table 2), we derived three binary variables representing social interaction types of interest: (1) all social interactions, (2) being groomed (with 1 assigned to ‘Groomed’ and ‘Grooming-groomed’) and (3) grooming (with 1 assigned to ‘Grooming’ and ‘Grooming-groomed’). For each interaction, we also included the matrilineal kinship (mother, sibling, same-matriline, non-kin/male) of the interaction partner.

### Inter-rater reliability

To ensure that observers recognised the individuals accurately, we conducted a training phase of several weeks, followed by a testing phase. Each test required the identification of an individual in three distinct situations: (1) at close range with high visibility, (2) at close range but with reduced visibility (either at height and/or obscured by dense foliage), and (3) at more than 10 metres. The tests were randomised both in terms of individuals and observation conditions. To validate individual recognition, an observer had to correctly identify the same individual across all three conditions in succession without error. If a mistake was made at any stage, the test for that individual restarted from the beginning (zero validated positions). This method, routinely used for all field assistants in the project, ensured a reliable and consistent identification of individuals before the collection of behavioural data.

We additionally assessed the reliability of behavioural coding by comparing the observations of the main coder and a secondary coder who blindly coded 20% of the dataset. Reliability was high across all categories with Cohen’s Kappa values of 0.82 for the global activities and 0.90 for the social activity types.

### Data Analyses

#### Proximity and social interactions with mother

To quantify the influence of maternal bonds on juvenile social development, we performed a two-part analysis. In the first analysis, we focused on the focal individual’s interactions with its mother compared to others. To do this, we got the proportion of a focal individual’s interactions per interacting individual, and we compared the proportion of interactions involving the mother versus the other individuals across all four interaction types (proximity, social interaction, grooming, and being groomed) and all age classes. In the second analysis, we extended this to investigate the role of maternal social rank on the proportion of interactions with the mothers and compared this proportion (interactions with their own mother) between individuals with a high-ranking mother to those with a low-ranking mother. We present the results for proximity and social interactions in the main manuscript and for both grooming interactions (being groomed and grooming) in the supplementary material as the latter were similar to the results of the social interactions.

#### Effect of socio-demographic factors on juvenile social profiles

To investigate how age and other factors affect individual social network profiles during juvenile development, we constructed two distinct sets of ego-centric multiplex networks for each of the two study groups. One set was based on proximity data, while the other was derived from social interaction data. Both multiplexes contained three layers, each corresponding to a different season. The association between individuals formed the intra-layer connections and the associations between the same individuals across layers formed the inter-layer connections. This approach allowed us to analyse the same set of individuals across multiple, temporally distinct networks simultaneously. We quantified each layer as undirected, weighted networks, where edges represent the frequency of interaction (weight) rather than simple presence/absence, and relationships are treated as reciprocal (undirected). Edge weights were normalised to sampling effort based on the type of dyad. For dyads consisting of a focal individual and a non-focal partner, the weight was calculated as the proportion of the focal individual’s samples in which the partner was in proximity or interacting with the focal. For dyads comprising two focal individuals, we adjusted for combined sampling effort by defining the weight as the total number of shared observations divided by the sum of samples collected for both individuals.

Using these networks, we quantified three key nodal properties that describe an individual’s social integration: multiplex degree, multiplex strength, and multiplex eigenvector centrality. Multiplex degree and multiplex strength represent the total number of connections and the sum of connection weights, respectively, for a given individual across all seasons [43]. In contrast, multiplex eigenvector centrality provides a more comprehensive measure of an individual’s overall importance or versatility across multiple social contexts [44]. This metric extends the traditional eigenvector centrality measure (which assesses the individual’s influence within a single network) by quantifying a node’s connectivity to other influential nodes across all layers. We calculated this value by first generating a supra-adjacency matrix that combines both intra-and inter-layer connections and then summing the eigenvector values of the leading eigenvector for each node across all layers.

#### Changes in network overlap across age

To investigate how network structure shifts with age, we performed two analyses using two distinct types of multiplex networks. For each analysis, we created a separate multiplex for each age class, season, and group, resulting in 18 unique networks (3 age classes, 3 seasons and 2 groups) for each type of multiplex network.

For our first analysis, we constructed multiplex networks with three layers: kinship, proximity, and social interaction. Each network contained only focal individuals of a specific age class and all individuals with whom they associated. The proximity and social interaction network layers were built as previously described with undirected weighted edges. To build the kinship layer, we first assigned a relationship score to each dyad: 3 for parent-offspring, 2 for siblings, 1 for other family members, and 0 for unrelated individuals. Since we only had maternal kinship information only this was taken into account in these networks. To ensure consistency with the proportional edge weights of the other layers, we normalised these scores to a scale from 0 to 1, where 1 represented the strongest relationship (parent-offspring) and 0 no relation. We included only individuals present in the proximity and social interaction network layers to ensure that all layers shared the same set of nodes.

In a second analysis, we constructed four-layer multiplex networks to compare specific social interactions and quantify reciprocity between types of social interactions. These network layers were kinship, proximity, grooming, and being groomed (referred to as groomed) layers. All four layers were directed, with edges moving outward from the focal individual. This approach allowed us to directly assess reciprocity: an edge from focal individual A to another individual B in both the grooming and being groomed networks indicates a reciprocal relationship, which would not be captured if edges were consistently directed from the giver to the receiver. To maintain consistency with this structure, we represented the symmetrical relationships between focal individuals in the kinship and proximity layers by including two oppositely directed edges with the same weight between two related or associated focal individuals.

Across all multiplexes, we quantified the similarity between layers using a weighted edge overlap metric. This metric is a single value, ranging from 0 (no overlap) to 1 (full similarity in weighted edges), that provides a comprehensive measure of how individuals’ connections align across different social contexts. We calculated this value by first taking the minimum edge weight between the same two nodes across the two layers, summing these minimum values across all node pairs, and then normalizing this sum by the total edge weights from both layers.

### Statistical analyses

We used generalised linear mixed-effects models (GLMMs) for all statistical analyses with different error structures depending on the analyses. For each analysis, we validated model assumptions, extracted estimated marginal means (EMMs) to visualise model predictions and performed post-hoc comparisons among predictor levels to identify significant differences.

To quantify the effect of maternal interactions and maternal rank, we fit separate GLMMs for each of the four interaction types (proximity, social interactions, grooming, and groomed). For the first analysis, the response variable was the proportion of interactions with each conspecific, and the fixed effects included an interaction term between interactor type (mother vs. others) and age class. This interaction was included because model comparisons using ANOVA revealed it significantly improved model fit. The models included random effects for season nested within group. The response variable was modelled using a beta distribution, which required a minor adjustment for values of 0 or 1, as described before. For the second analysis, to test the effect of maternal rank on the proportion of interactions with the mothers, we used the same model structure but with an interaction term between interactor type (focal with high-ranking mother vs. focal with low-ranking mother) and age class. In this case, the interaction term was not significant, so we used a simpler model with only the main effects of interactor type and age class.

To determine the drivers of individual network properties, we fit separate GLMMs for each network type (proximity and social interactions) and each nodal property (multiplex degree, strength, and eigenvector centrality). In all models, the response variable was the specific nodal property, while fixed effects included age class (3 levels: [<1yo], [1yo] and [2yo]), sex (2 levels, male and female), maternal rank (2 levels: high and low) and group (2 levels: AK and KB). The model error structures were chosen to match the response variable type: a Poisson distribution for multiplex degree, a Gaussian distribution for multiplex strength, and a beta distribution for eigenvector centrality. Since the beta distribution is defined on the open interval (0,1), we applied a minor adjustment to any values of 0 or 1 to ensure model compatibility [45].

To quantify age related changes in edge overlap between pairs of network layers in the multiplexes, we fit separate GLMMs for each pair of layers (kinship-proximity, kinship-social, proximity-social, kinship-grooming, kinship-groomed, proximity-grooming, proximity-groomed, grooming-groomed). In all models, the response variable was the edge overlap value, the fixed effect was the age class (3 levels: [<1yo], [1yo] and [2yo]), and the random error structure was season nested within group with a beta distribution as the model error structure.

Multiplex networks were created and analysed using the muxViz package in R. Statistical analyses were performed using the glmmTMB, modelbased, performance, DHARMa and emmeans packages in R [46–48]. Plots were created using ggplot, ggh4x, ggtext, cowplot and multcomp View packages in R [49–53].

## Results

### 1. Proximity and social interactions with mother

Juveniles spent more time in proximity to their mothers compared to other partners before 2yo (Fig. 1A, Table S1; in [<1yo]: difference between others and maternal proportion = -0.26; CI = [-0.32, -0.19], *p* < 0.001; in [1yo]: difference = -0.06; CI = [-0.10, -0.02], *p* = 0.006; in [2yo]: difference = 0.02; CI = [-0.01, 0.06], *p* = 0.255) and also engaged in more social interactions with their mothers before they turn 2-year old (Fig. 1A, Table S1; in [<1yo]: difference = -0.48; CI = [-0.61, - 0.35], *p* < 0.001; in [1yo]: difference = -0.15; CI = [-0.25, -0.06], *p* = 0.001; in [2yo]: difference = - 0.05; CI = [-0.14, 0.04], *p* = 0.283). With age, the time spent in proximity and in social interactions with the mothers decreased (Table S1; p < 0.05 for comparisons between all age classes in proximity and for comparisons between [<1yo]-[1yo] and [<1yo]-[2yo] in social interactions).

**Figure 1:**
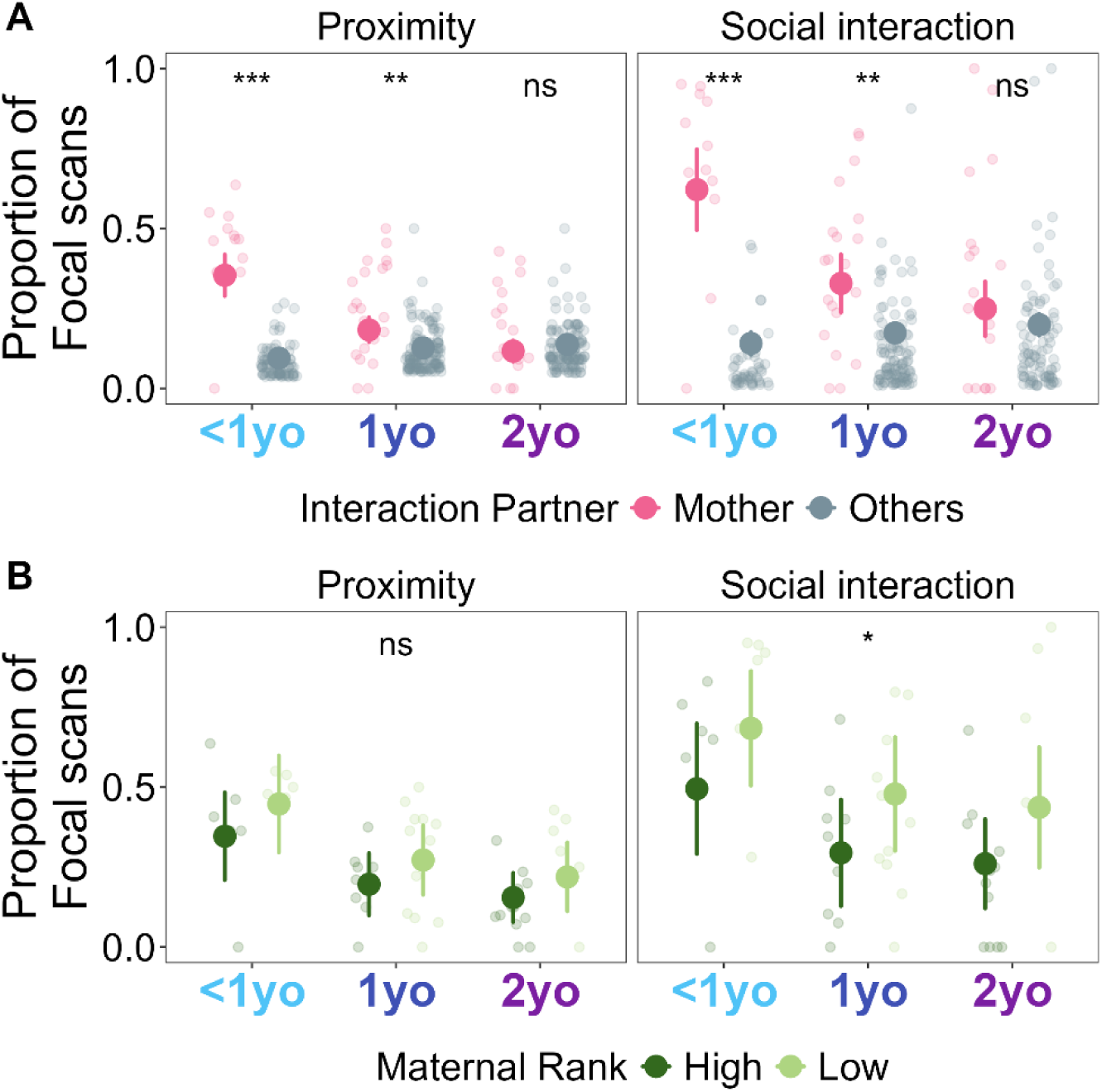
Age-related changes in grooming received from, and given to, the mother by juveniles. A) Proportion of scan samples that focal individuals engaged in social interactions with their mothers and other conspecifics across age classes. Smaller circles represent data from one interaction dyad (focal - conspecific) while larger circles represent the estimated marginal means with the 95% confidence interval represented by error bars. B) Proportion of scan samples that focal individuals with high-ranking and low-ranking mothers engaged in social interactions with their mothers across age classes. Smaller circles represent interaction from one focal-maternal pair while larger circles represent the estimated marginal means with the 95% confidence interval represented by error bars. Asterisks indicate statistically significant differences based on post hoc comparisons (ns = p > 0.05, * = 0.05 > p > 0.01; ** = 0.01 > p > 0.001; *** = p < 0.001).

Maternal rank did not influence the time spent by juveniles in proximity of their mother (Fig. 1B, Table S2; Low vs High rank: difference = 0.08, CI = [-0.02, 0.18], *p* = 0.103). However, juveniles with low-ranking mothers spent a greater proportion of time in social interactions with their mothers (Fig. 1B, Table S2; Low vs High rank: difference = 0.18, CI = [0.04, 0.32], *p* = 0.01).

### 2. Effect of socio-demographic factors on juvenile social profiles

Only age and maternal rank significantly affected the multiplex centrality metrics in the proximity network (Fig. 2A, Table S3). Specifically, the bond strength (multiplex strength) decreased significantly between [<1yo] and the older ones (compared to [1yo]: difference= 0.65, CI = [0.12, 1.17], *p* = 0.010; compared to [2yo]: difference= 0.79, CI = [0.36, 1.22], *p* = 0.001). Individuals with high-ranking mothers showed higher diversity of partners (multiplex degree), stronger bonds (multiplex strength) and greater connections to influential partners (that is, partners who were themselves well-connected and influential within the broader network; multiplex eigenvector centrality) compared to those with low-ranking mothers (multiplex degree: = -0.23, CI = [-0.42, - 0.05], *p* = 0.015; multiplex strength: difference = -0.42, CI = [-0.66, -0.17], *p* = 0.003; multiplex eigenvector centrality: difference =-1.47, CI = [-2.23, -0.71], *p* < 0.001).

**Figure 2:**
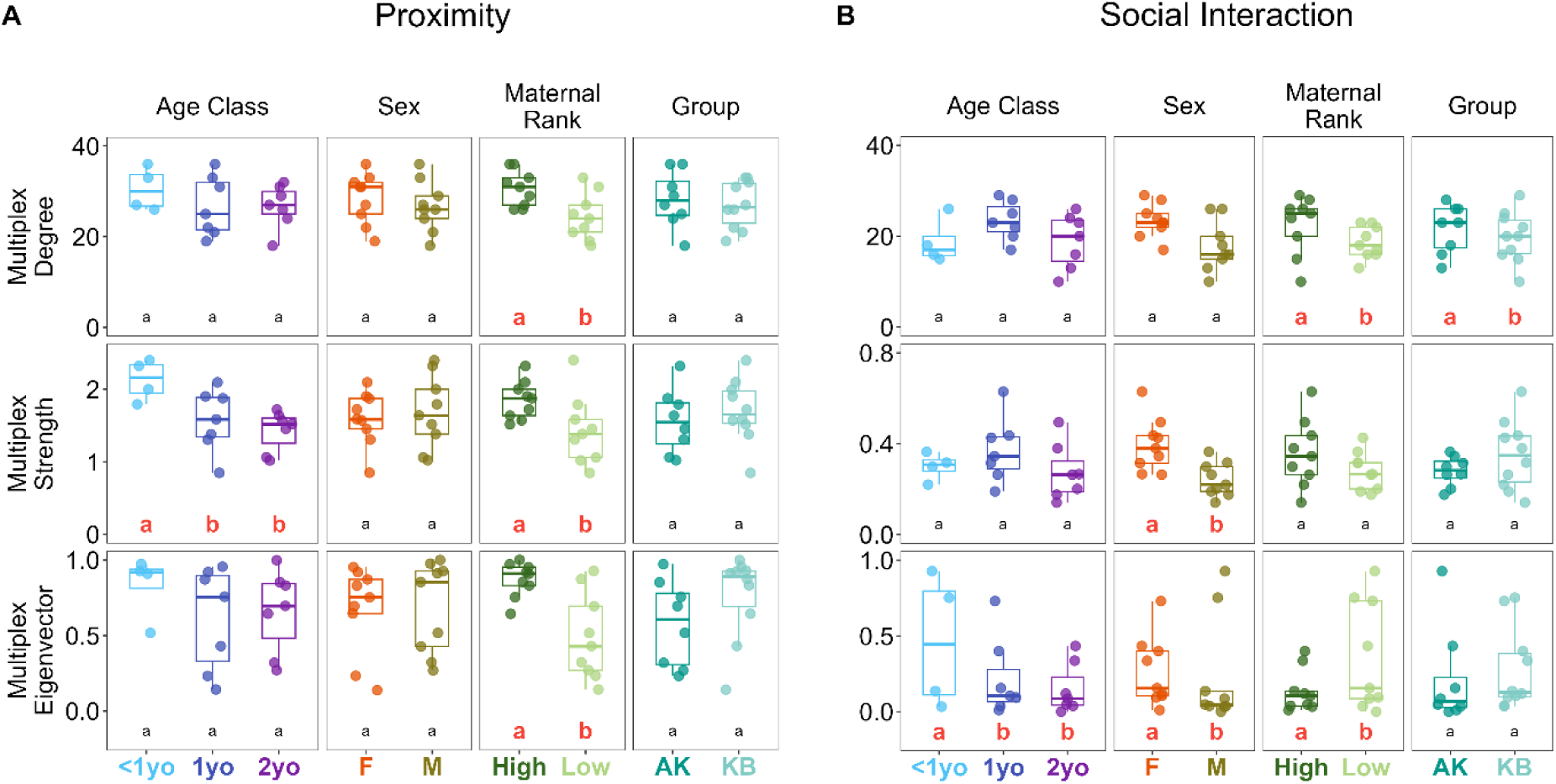
Effects of socio-demographic factors on juvenile multilayer centrality metrics in A) proximity and B) social interaction networks. Each circle represents data from one focal individual, and the boxplot represents the distribution for each category, with the median given by the horizontal line, and the box representing the interquartile range. Letters indicate statistically significant differences based on post hoc comparisons within each socio-demographic category and comparisons involving significant differences (p < 0.05) are highlighted with larger red font.

All the tested factors affected the multiplex centrality metrics in the social interaction networks (Fig. 2B, Table S4). Specifically, the connection to influential partners decreased significantly between [<1yo] and older age classes (multiplex eigenvector centrality in comparison to [1yo]: difference = 2.65, CI = [0.82, 4.47], *p* = 0.002; to [2yo]: difference = 1.96, CI = [0.49, 3.42], *p* = 0.003). Juveniles with high-ranking mothers exhibited higher partner diversity but lower influential connections within the network compared to those with low-ranking mothers (multiplex degree: difference = -0.23, CI = [-0.44, -0.02], *p* = 0.035; multiplex eigenvector centrality: difference = 1.42, CI = [0.51, 2.33], *p* = 0.002). Juvenile females showed stronger bonds and higher connections to influential partners within the network than juvenile males (multiplex strength: difference between females and males = 0.18, CI = [0.06, 0.29], *p* = 0.005; multiplex eigenvector centrality: difference = 1.55, CI = [0.41, 2.77], *p* = 0.008). And at the group level, AK juveniles showed higher diversity of social partners than KB ones (multiplex degree: difference between AK and KB = -0.23, 95% CI = [-0.45, 0.01], *p* = 0.036).

### 3. Changes in network overlap across age

#### 3.1. Similarity across proximity, social interactions, and kinship networks

The edge overlap between kinship and proximity networks significantly decreased from [<1yo] to [1yo] and then stabilised further (Fig. 3, Table S5; [1yo] vs [<1yo]: difference = -0.07, CI = [-0.12, - 0.02], *p* = 0.005; [2yo] vs [<1yo]: difference = -0.11, CI = [-0.15, -0.06], *p <* 0.001; [2yo] vs [1yo]: difference = -0.04, CI = [-0.08, 0.01], *p =* 0.123).

**Figure 3:**
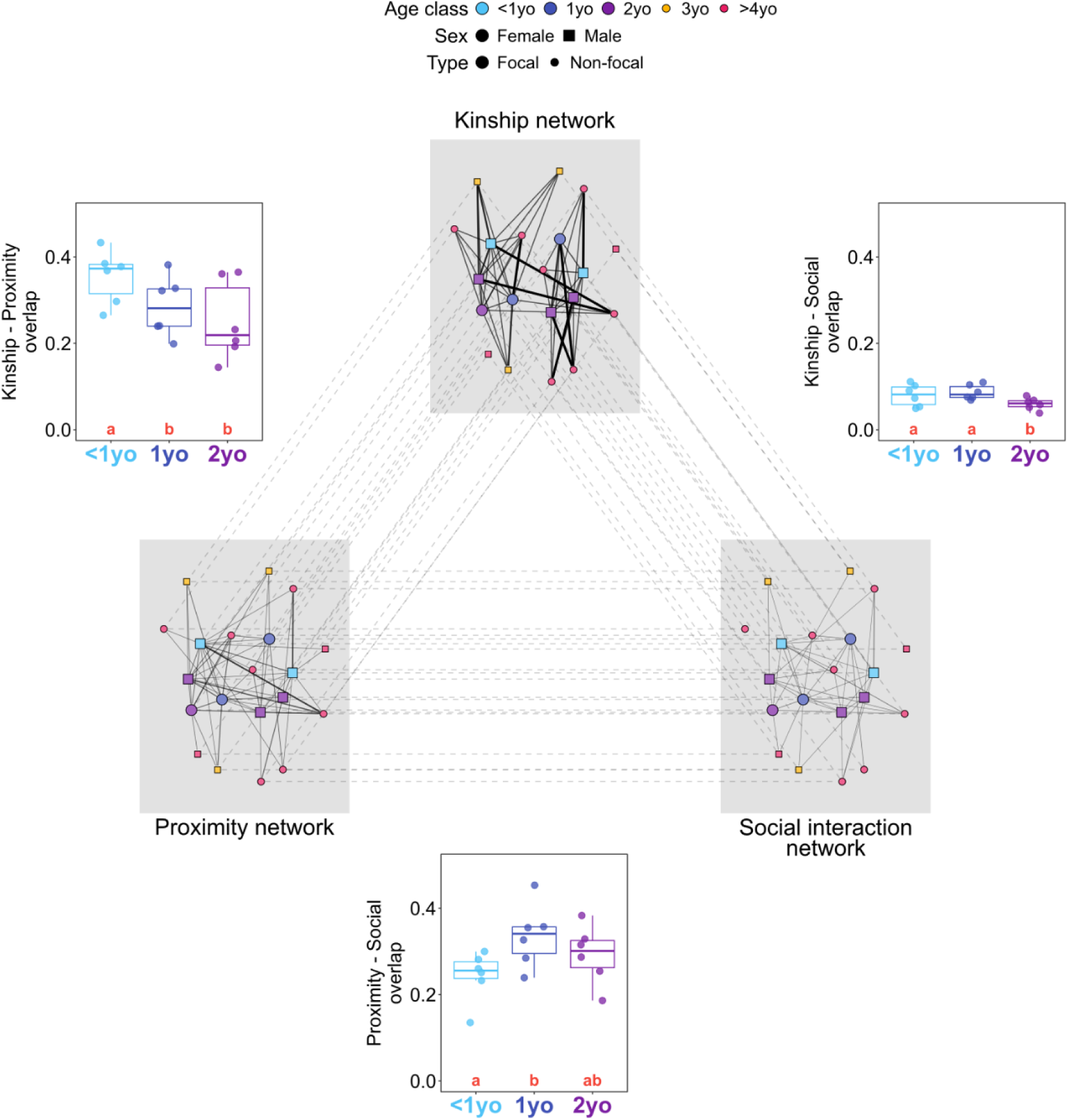
Edge overlap between proximity, kinship and social interaction layers across age classes in multiplex networks. The networks in grey in the centre represent data from group AK in summer. Each node represents an individual in the group with the node colour representing their age, the shape their sex and the size representing whether they are a focal juvenile or a non-focal conspecific. Intra-layer edges connect dyads while inter-layer edges connect the same individuals. The graphs between each pair of layers represent the edge overlap between these layers over age. Each circle represents data from one group and season. Boxplots represent the distribution for each age class, with the median given by the horizontal line, and the box representing the interquartile range. Letters indicate statistically significant differences across age classes based on post hoc comparisons and comparisons involving significant differences (*p* < 0.05) are highlighted with larger red font.

The edge overlap between kinship and social interaction networks was stable from [<1yo] to [1yo] and then significantly decreased further (Fig. 3, Table S5; [1yo] vs [<1yo]: difference = 0.01, CI = [-0.01, 0.03], *p* = 0.361; [2yo] vs [<1yo]: difference = -0.02, CI = [-0.04, -0.01], *p =* 0.022; [2yo] vs [1yo]: difference = -0.03, CI = [-0.04, -0.01], *p* = 0.001).

On the other hand, the edge overlap between proximity and social interaction networks significantly increased from [<1yo] to [1yo] and then stabilised further (Fig. 3, Table S5; [1yo] vs [<1yo]: difference = 0.09, CI = [0.04, 0.15], *p* = 0.001; [2yo] vs [<1yo]: difference = 0.04, CI = [-0.01, 0.10], *p =* 0.081; [2yo] vs [1yo]: difference = -0.04, CI = [-0.10, 0.01], *p* = 0.128).

#### 3.2. Similarity across proximity, kinship and allogrooming networks

##### 3.2.1. Similarity between kinship and allogrooming networks (i.e., grooming and groomed)

The edge overlap between kinship and grooming networks significantly increased from [<1yo] to [1yo] and then significantly decreased further (Fig. 4A, Table S6; [1yo] vs [<1yo]: difference = 0.13; CI = [0.10, 0.16], *p* < 0.001; [2yo] vs [<1yo]: difference = 0.08; CI = [0.06, 0.10], *p* < 0.001; [2yo] vs [1yo]: difference = -0.05; CI = [-0.08, -0.02], *p* = 0.004). The edge overlap between kinship and being groomed networks significantly increased from [<1yo] to [1yo] and then stabilised further (Fig. 4A, Table S6; [1yo] vs [<1yo]: difference = 0.04; CI = [0.01, 0.08], *p* = 0.007; [2yo] vs [<1yo]: difference = 0.03; CI = [0, 0.06], *p* = 0.049; [2yo] vs [1yo]: difference = -0.01; CI = [-0.05, 0.02], *p* = 0.444).

**Figure 4:**
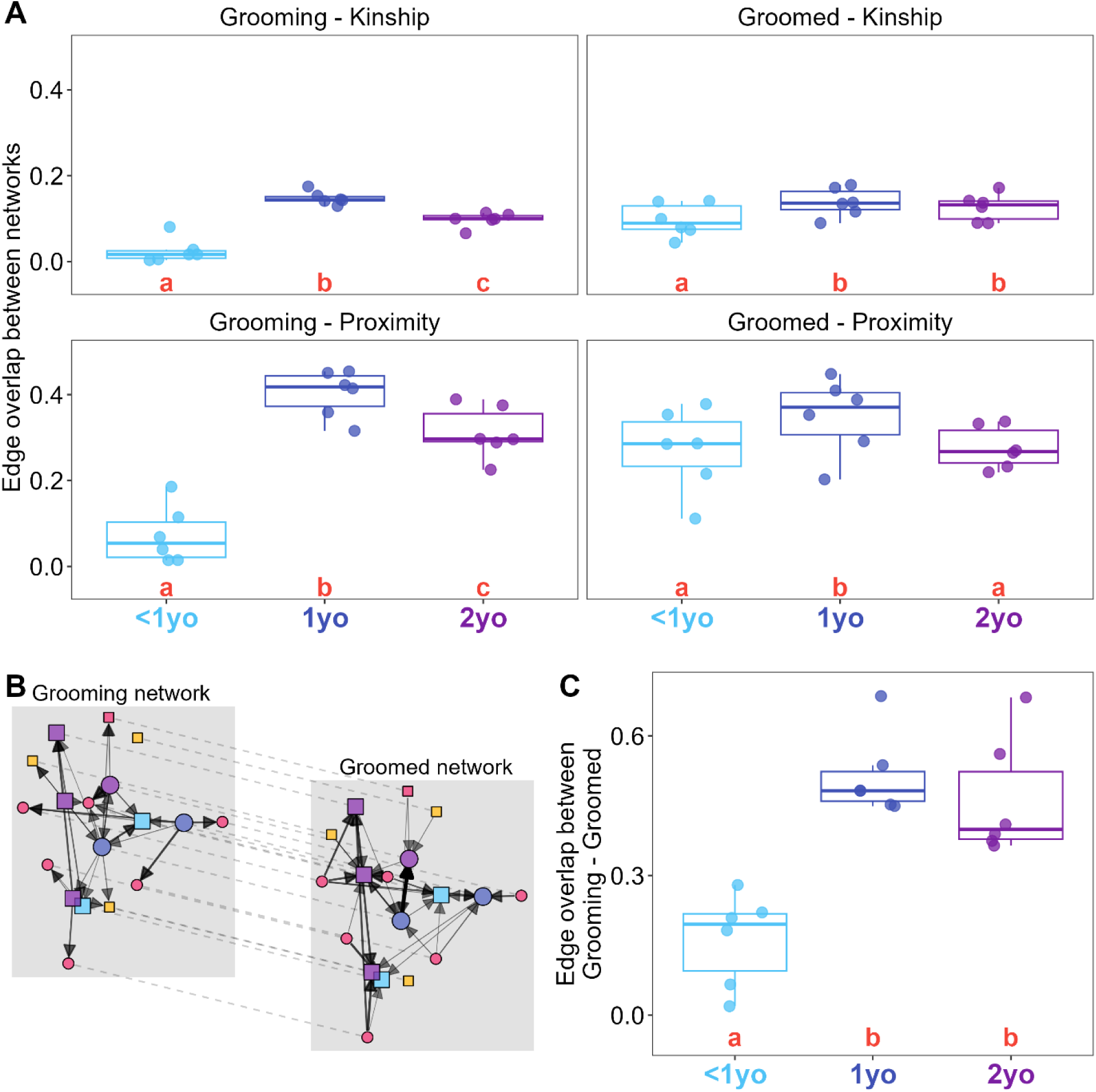
Edge overlap between proximity, kinship and grooming-groomed networks across age classes in multiplex networks. A) The edge overlap between grooming and being groomed networks with kinship and proximity networks. Each circle represents data from one group and season. Boxplots represent the distribution for each age class, with the median given by the horizontal line, and the box representing the interquartile range. Letters indicate statistically significant differences across age classes based on post hoc comparisons and comparisons involving significant differences (p < 0.05) are highlighted with larger red font. B) Representative plot of grooming and groomed network layers from the multiplex from group AK in summer. The nodes represent individuals in the group, and their colour, shape and size follow the same pattern as in Fig. 3. The edges are directed, with the arrows going from the individual performing the grooming to the individual being groomed and their size is weighted based on the proportion of grooming interactions. C) Edge overlap between grooming and groomed networks across age class. The circles, boxplots and significance letters are the same as in A.

##### 3.2.2. Similarity between proximity and allogrooming networks (i.e., grooming and groomed)

The edge overlap between proximity and grooming networks <u>s</u>ignificantly increased from [<1yo] to [1yo] and then significantly decreased further (Fig. 4A, Table S6; [1yo] vs [<1yo]: difference = 0.34; CI = [0.27, 0.41], p < 0.001; [2yo] vs [<1yo]: difference = 0.25; CI = [0.19, 0.32], p < 0.001; [2yo] vs [1yo]: difference = -0.09; CI = [-0.18, -0.004], p = 0.042). The edge overlap between proximity and groomed networks significantly increased from [<1yo] to [1yo] and then significantly decreased to initial levels (Fig. 4A, Table S6; [1yo] vs [<1yo]: difference = 0.08; CI = [0.02, 0.14], *p* = 0.012; [2yo] vs [<1yo]: difference = 0.01; CI = [-0.05, 0.07], *p* = 0.816; [2yo] vs [1yo]: difference = -0.07; CI = [-0.14, -0.01], *p* = 0.024).

##### 3.2.3. Similarity grooming and groomed networks

The edge overlap between grooming and groomed networks significantly increased from [<1yo] to [1yo] and then stabilised (Fig. 4B and C, Table S6; [1yo] vs [<1yo]: difference = 0.37; CI = [0.26, 0.48], *p* < 0.001; [2yo] vs [<1yo]: difference = 0.32; CI = [0.21, 0.43], *p* < 0.001; [2yo] vs [1yo]: difference = -0.05; CI = [-0.18, 0.08], *p* = 0.456).

## Discussion

In this study, we investigated juvenile social development by examining the temporal dynamics of maternal independence, associated changes in juvenile centrality in their ego networks and the structural alignment between kinship, proximity and social interaction networks. Our results show that the social development of juvenile vervet monkeys is a transitory phase of social reorganisation, driven primarily by its growing independence from its mother. This transition is marked by a shift in the drivers of association, where social interactions evolve from being structured by matrilineal kinship to being increasingly shaped by spatial proximity. Our analysis further reveals how this developmental trajectory is shaped by inherited social status, through the effects of maternal rank, and the precocial emergence of sex-specific social roles.

The changing nature of the mother-offspring bond is the primary driver of a juvenile vervet’s early social development. As expected in a matrilineal society, our results show that newborns are clearly first oriented towards their mothers in both their spatial proximity and direct social interactions [54]. However, we observed a progressive decline in this maternal focus over time. This gradual weakening of the maternal focus can arise from mothers reducing contact and increasing rejection behaviours, while other group members (i.e., grandmothers and young females) begin to provide care [54–58]. Juveniles also start to direct more attention toward matrilineal kin, peers, and high-ranking adults who serve as social models [27,29,59]. By around two years of age, juveniles no longer show a strong preference for their mother over other group members in our study, consistent with developmental patterns found in other matrilineal species as macaques [60]. Notably, we find that this decline is more pronounced for social interactions than for proximity, suggesting a redirection of social investment rather than merely a change in positioning patterns.

This developmental trajectory is further shaped by the mother’s social standing in a targeted manner. While maternal rank did not influence the time mother-offspring dyads spent near each other, it significantly affected their social interactions. Low-ranking mothers engaged in a significantly higher proportion of social interactions with their offspring than high-ranking mothers, a pattern also observed in rhesus macaques and Japanese macaques, where subordinate females maintained closer contact and longer suckling bouts [61–63]. This disparity likely reflects a compensatory social strategy with two complementary functions. First, it may help offset the limited social capital available to the offspring of low-ranking females. Second, it may provide a richer learning context, enabling low-ranking mothers to better equip their offspring with the social skills needed to navigate a challenging social landscape [27,29]. In this way, subordinate mothers strategically shape their offspring’s early social experience, mitigating the constraints of their inherited social status.

These maternal-driven changes shape a developmental process characterised not by social expansion, but by social reorganisation (Fig. 5). Contrary to the expectation that independence would broaden a juvenile’s network, the number of interaction partners (network degree) remained remarkably stable with age in both proximity and social interaction networks. However, this overarching stability masked subtle but significant shifts. In proximity networks, connection strength weakened, mirroring findings by Vilette and colleagues [21]. In social interaction networks, we saw a reduction in ties to influential partners (eigenvector centrality), a decline likely initiated by the waning influence of the mother as the juvenile’s primary social anchor. This latter finding diverges from Vilette and colleagues [21], who found that juveniles strengthened their grooming connections, suggesting that the developmental trajectory of social bonds depends on the specific type of interaction being measured. Overall, this pattern of reorganisation strongly indicates that as juveniles mature, they channel their growing independence not into social expansion, but into fitness related activities like foraging and navigation [64].

**Figure 5:**
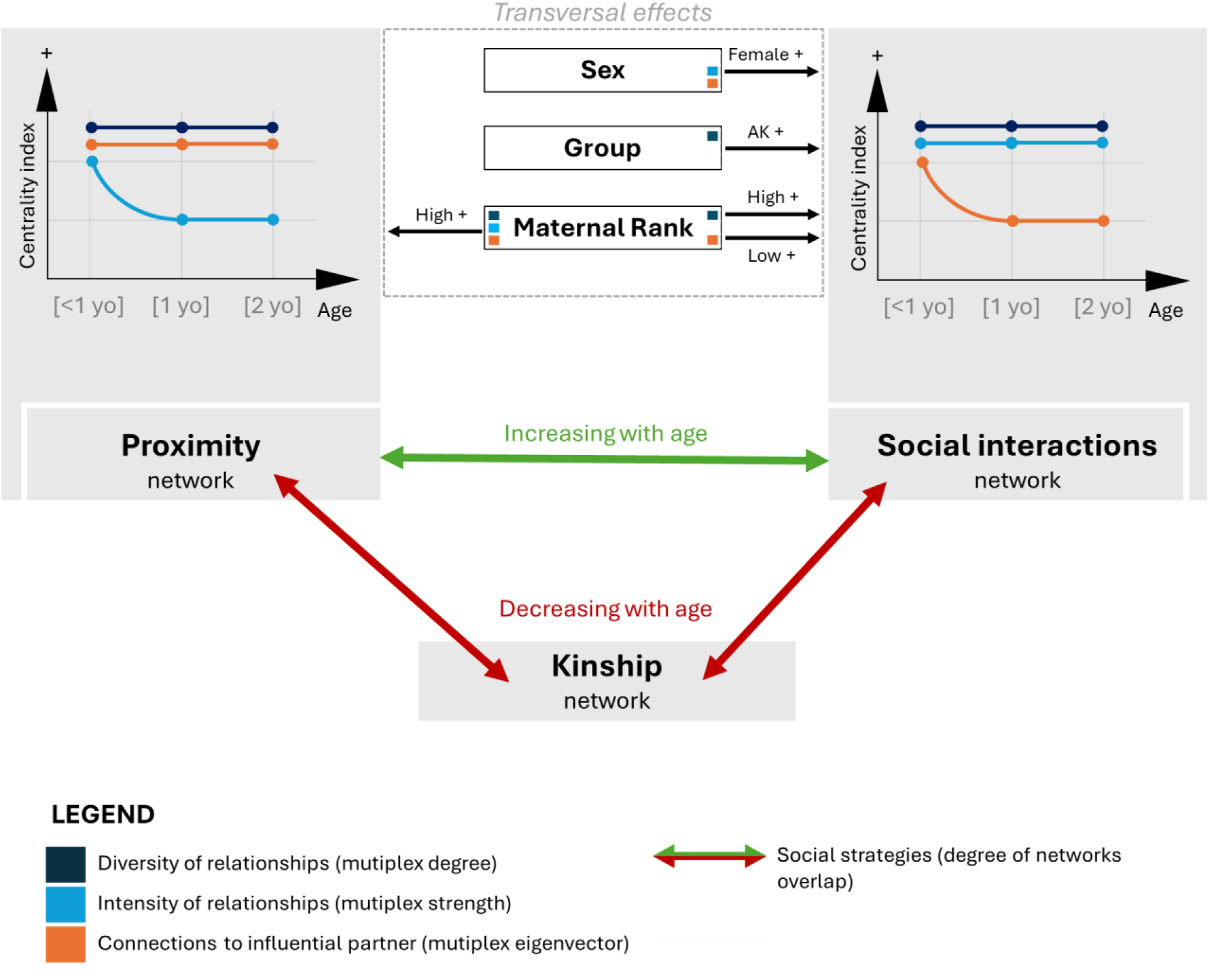
Summary of developmental and transversal effects on multilayer social network structure. The upper panels show significant effects of age, maternal rank, sex and group on node centrality within each network. The lower arrows summarise age-related changes in the degree of overlap between proximity, social interaction and kinship networks. The arrows indicate significant positive (green) or negative (red) effects of the factors on the networks, and then, significant effect of age on the degree of overlap between networks.

Beyond the general developmental pattern of network reorganisation, our analysis revealed that variation in juveniles’ social networks was driven far more by sex than by social group (Fig. 5). Although the groups differed in their social dynamics [28,65,66], this variation had only a negligible effect on juveniles’ network centrality. We also found a group effect only on the multiplex degree in social interaction networks. Given that group sizes were comparable (AK = 23; KB = 26), this difference is likely related to qualitative differences in social structure, specifically the higher social tolerance in AK [64,66]. In contrast, sex had a strong influence: males and females occupied comparable positions in proximity networks, however juvenile females were significantly more central in social interaction networks (as found in [24] in grooming networks). This finding, consistent with Vilette and colleagues [21], is particularly remarkable given that these sex-specific patterns emerge long before major life-history events such as male dispersal [65,67] and even before the development of secondary sexual characteristics (e.g., body size and genital allometry; [68]). Our results indicate that the foundations of adult social strategies are laid in juvenescence, as females invest in the affiliative bonds supporting lifelong philopatry, whereas males likely prioritise play and other skills relevant to future dispersal [25].

Interestingly, our results indicate that juvenile social development may partly arise from two complementary patterns in social strategies: an increasing alignment between proximity and interaction networks, and a parallel decline in kin-based associations (Fig. 5). As juveniles gain independence from their mothers, their relationships become less centred on kin and progressively stabilise across a broader range of group members, consistent with a general trajectory of social independence. On the other hand, with age, the individuals increasingly spend time near those with whom they interact most, indicating a tighter coupling between spatial association and social engagement (i.e., increasing spatial-based strategy). Intriguingly, these developmental dynamics do not follow a simple linear progression. We observed a temporary peak in proximity-interaction overlap around the age of one year old, followed by a slight decline at two years, approaching the levels observed before one year. Thus, the transition toward social autonomy is not gradual but marked by periods of adjustment and recalibration [26].

Furthermore, this non-linear developmental pattern is most evident in grooming behaviour. We observed a clear peak around one year of age in how grooming aligns with proximity and kinship, followed by a decline by two years. The transition in grooming interactions also appears to set the stage for more reciprocal relationships: the overlap between whom a juvenile grooms and by whom it is groomed increases sharply around one year and remains high at two years. Thus, it seems likely that, once established, reciprocal exchanges become a lasting feature of juveniles’ social relationships. These changes coincide with a reduction in maternal involvement. Early grooming in juveniles is mostly provided by mothers, but around one year this imbalance fades as juveniles begin to engage in more balanced exchanges with others (Fig. S1, Table S2). Notably, juveniles of low-ranking mothers both gave and received more grooming, supporting the idea that these mothers invest extra social effort to help their offspring build relationships and navigate their social environment (Fig. S1, Table S2).

## Conclusion

By employing a multilayer network approach, our study shows that vervet social development is characterised by a qualitative, not quantitative, shift. Instead of an expansion of the juvenile social world, there is a shift in the factors structuring social association, a process associated with the weakening mother-offspring bond. Our analysis highlights the specific dynamics of this transition: proximity-based interactions peak around the one-year mark, whereas kin-based interactions, rather than just declining, are refocused, with grooming of kin increasing with age. Ultimately, our work shows how using a multilayer framework to quantify social development moves beyond simple metrics of sociality to reveal the complex interplay between inherited status and social relationships in juvenile primates. Vervet monkeys thus provide a powerful model for studying the systemic development of social behaviour (e.g., the non-linear developmental modal of Representational Redescription [69]) within a comparative perspective.

## Acknowledgments

This work was supported by a Fyssen Foundation postdoctoral fellowship to MD. At the time of writing, CC was supported by the CNRS and by the Interdisciplinary Thematic Institute NeuroStra (ITI 2021–2028 programme of the University of Strasbourg, CNRS, Inserm, IdEx Unistra ANR-10-IDEX-0002) under the framework of the French Programme “Investments for the Future”. The field work of this project and EvdW’s salary were supported by the European Research Council under the European Union’s Horizon 2020 research and innovation programme for the ERC ‘KNOWLEDGE MOVES’ starting Grant to EvdW (grant agreement No. 949379). EAG was funded by the Department of Ecology and Evolution at the University of Lausanne. We thank the Simian Laboratory Europe (Silabe, University of Strasbourg, France) for welcoming MD, CC and FB during the analysis and writing of this research. We are grateful to Ismaël Kerdli for his contribution to the behavioural coding of the videos. We also thank the field assistants and IVP managers for their invaluable support during data collection.

