## Supplementary Information for "From kin to spatial proximity conspecifics: social network reorganisation in developing juvenile vervet monkeys"

### Results

#### 1. Maternal vs non-maternal partners

**Table S1. Estimated differences between maternal and non-maternal partners across age groups.**

| Predictor | Pred_cat | Level1 | Level2 | Diff. | SE | CI_low | CI_high | z | p | |
|---|---|---|---|---|---|---|---|---|---|---|
| Proximity | | | | | | | | | | |
| AgeR | <1yo | others | maternal | -0,26 | 0,03 | -0,32 | -0,19 | -7,70 | **p<0,001** | *** |
| | 1yo | others | maternal | -0,06 | 0,02 | -0,10 | -0,02 | -2,76 | **0,006** | ** |
| | 2yo | others | maternal | 0,02 | 0,02 | -0,01 | 0,06 | 1,14 | 0,255 | |
| Partner | maternal | 1yo | <1yo | -0,17 | 0,04 | -0,25 | -0,10 | -4,48 | **p<0,001** | *** |
| | | 2yo | <1yo | -0,24 | 0,04 | -0,31 | -0,16 | -6,45 | **p<0,001** | *** |
| | | 2yo | 1yo | -0,07 | 0,03 | -0,12 | -0,02 | -2,59 | **0,010** | * |
| | others | 1yo | <1yo | 0,03 | 0,01 | 0,01 | 0,05 | 3,43 | **0,001** | ** |
| | | 2yo | <1yo | 0,04 | 0,01 | 0,02 | 0,06 | 4,53 | **p<0,001** | *** |
| | | 2yo | 1yo | 0,01 | 0,01 | -0,01 | 0,03 | 1,18 | 0,238 | |
| Social interactions | | | | | | | | | | |
| AgeR | <1yo | others | maternal | -0,48 | 0,07 | -0,61 | -0,35 | -7,14 | **p<0,001** | *** |
| | 1yo | others | maternal | -0,15 | 0,05 | -0,25 | -0,06 | -3,21 | **0,001** | ** |
| | 2yo | others | maternal | -0,05 | 0,05 | -0,14 | 0,04 | -1,07 | 0,283 | |
| Partner | maternal | 1yo | <1yo | -0,29 | 0,08 | -0,45 | -0,14 | -3,68 | **p<0,001** | *** |
| | | 2yo | <1yo | -0,37 | 0,08 | -0,53 | -0,22 | -4,77 | **p<0,001** | *** |
| | | 2yo | 1yo | -0,08 | 0,06 | -0,20 | 0,04 | -1,25 | 0,210 | |
| | others | 1yo | <1yo | 0,03 | 0,02 | -0,01 | 0,08 | 1,49 | 0,137 | |
| | | 2yo | <1yo | 0,06 | 0,02 | 0,01 | 0,11 | 2,47 | **0,014** | * |
| | | 2yo | 1yo | 0,03 | 0,02 | -0,02 | 0,07 | 1,21 | 0,228 | |
| Grooming | | | | | | | | | | |
| AgeR | <1yo | others | maternal | -0,05 | 0,05 | -0,14 | 0,05 | -1,01 | 0,311 | |
| | 1yo | others | maternal | -0,05 | 0,05 | -0,14 | 0,05 | -0,98 | 0,326 | |
| | 2yo | others | maternal | -0,05 | 0,05 | -0,15 | 0,05 | -0,99 | 0,321 | |
| Partner | maternal | 1yo | <1yo | -0,05 | 0,07 | -0,20 | 0,09 | -0,71 | 0,480 | |
| | | 2yo | <1yo | 0,01 | 0,08 | -0,15 | 0,16 | 0,07 | 0,944 | |

| | | | | | | | | | | |
|---|---|---|---|---|---|---|---|---|---|---|
| | | 2yo | 1yo | 0,06 | 0,04 | -0,03 | 0,15 | 1,31 | 0,190 | |
| | others | 1yo | <1yo | -0,05 | 0,07 | -0,19 | 0,09 | -0,69 | 0,488 | |
| | | 2yo | <1yo | 0,01 | 0,07 | -0,14 | 0,15 | 0,07 | 0,944 | |
| | | 2yo | 1yo | 0,05 | 0,04 | -0,03 | 0,14 | 1,30 | 0,194 | |
| Groomed | | | | | | | | | | |
| AgeR | <1yo | others | maternal | -0,49 | 0,08 | -0,64 | -0,33 | -6,05 | **p<0,001** | *** |
| | 1yo | others | maternal | -0,12 | 0,07 | -0,26 | 0,01 | -1,79 | 0,073 | |
| | 2yo | others | maternal | 0,10 | 0,07 | -0,03 | 0,23 | 1,52 | 0,129 | |
| Partner | maternal | 1yo | <1yo | -0,29 | 0,09 | -0,47 | -0,11 | -3,13 | **0,002** | ** |
| | | 2yo | <1yo | -0,43 | 0,09 | -0,60 | -0,26 | -4,92 | **p<0,001** | *** |
| | | 2yo | 1yo | -0,14 | 0,08 | -0,30 | 0,01 | -1,79 | 0,074 | |
| | others | 1yo | <1yo | 0,08 | 0,05 | -0,02 | 0,17 | 1,52 | 0,129 | |
| | | 2yo | <1yo | 0,15 | 0,06 | 0,05 | 0,26 | 2,78 | **0,005** | ** |
| | | 2yo | 1yo | 0,08 | 0,05 | -0,02 | 0,17 | 1,57 | 0,115 | |

**Table S2. Estimated differences in proximity and social interactions between High-ranking and Low-ranking partners across age groups.**

| Predictor | Pred_cat | Level1 | Level2 | Diff. | CI_low | CI_high | SE | z | p | |
|---|---|---|---|---|---|---|---|---|---|---|
| Proximity | MothR | Low | High | 0,08 | -0,02 | 0,18 | 0,05 | 1,63 | 0,103 | |
| Social | MothR | Low | High | 0,18 | 0,04 | 0,32 | 0,07 | 2,57 | **0,010** | * |
| Grooming | MothR | Low | High | 0,19 | 0,01 | 0,37 | 0,09 | 2,08 | **0,037** | * |
| Groomed | MothR | Low | High | 0,17 | 0,02 | 0,32 | 0,08 | 2,26 | **0,024** | * |

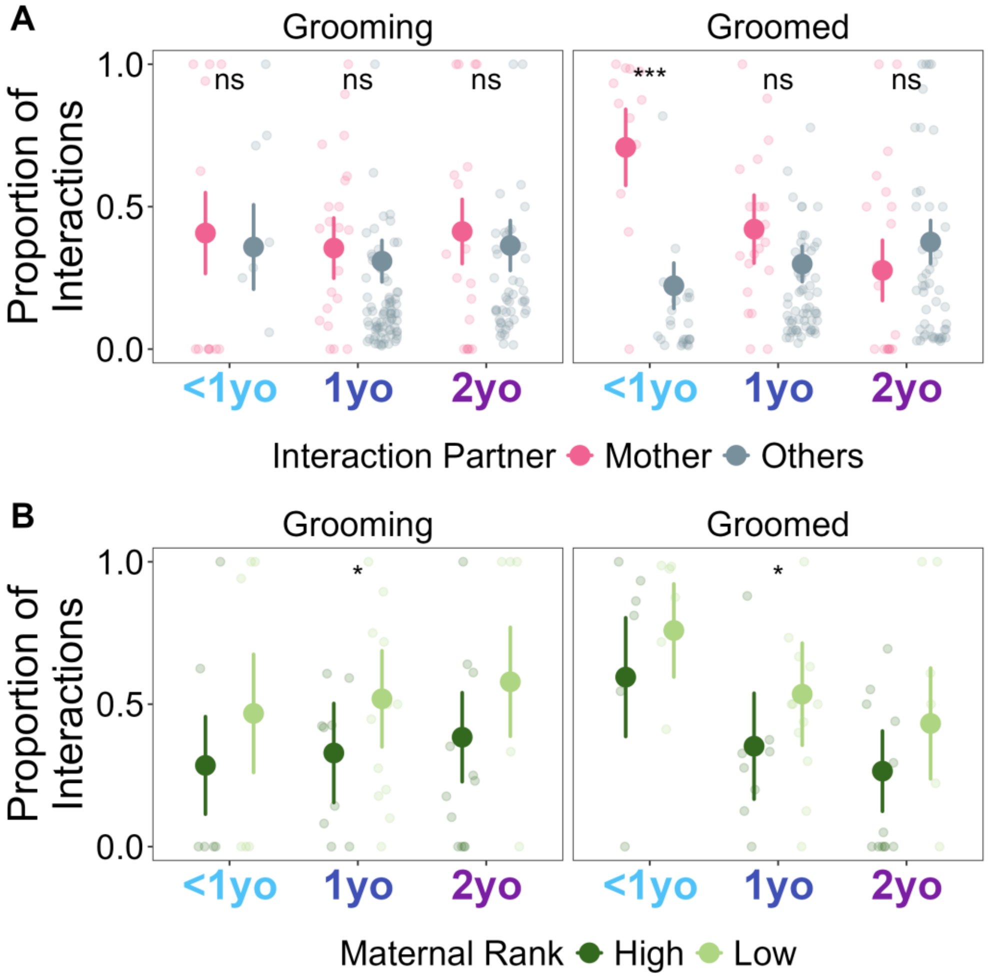


**Figure S1. Age-related changes in grooming given and received by the mother, in juveniles.** A) Proportion of scan samples that focal individuals spent in proximity with and engaged in social interactions with their mothers and other conspecifics across age classes. Smaller circles represent data from one interaction dyad (focal - conspecific) while larger circles represent the estimated marginal means with the 95% confidence interval represented by error bars. B) Proportion of scan samples that focal individuals with high-ranking and low-ranking mothers spent in proximity with and engaged in social interactions with their mothers across age classes. Smaller circles represent interaction from one focal-maternal pair while larger circles represent the estimated marginal means with the 95% confidence interval represented by error bars. Asterisks indicate statistically significant differences based on post hoc comparisons (ns = $p > 0.05$, * = $0.05 > p > 0.01$; ** = $0.01 > p > 0.001$; *** = $p < 0.001$).

#### 2. Effect of structural factors (i.e., age, sex, maternal rank and group)

**Table S3. Effects of age, sex, maternal rank (MatR) and social group on multiplex metrics (i.e., multiplex degree, strength and eigenvector) of the *1 meter-proximity* network.**

| | Pred. | Level1 | Level2 | Diff. | CI_low | CI_high | SE | t/z | p | |
|---|---|---|---|---|---|---|---|---|---|---|
| Multip. | Age | <1yo | 1yo | 0,26 | -0,13 | 0,66 | 0,16 | 1,608 | 0,216 | |
| Degree | | <1yo | 2yo | 0,24 | -0,07 | 0,55 | 0,13 | 1,834 | 0,200 | |
| | | 1yo | 2yo | -0,02 | -0,31 | 0,26 | 0,12 | -0,195 | 0,846 | |
| | Sex | Female | Male | 0,20 | -0,06 | 0,45 | 0,13 | 1,527 | 0,127 | |
| | MatR | Low | High | -0,23 | -0,42 | -0,05 | 0,09 | -2,435 | **0,015** | * |
| | Group | KB | AK | -0,12 | -0,31 | 0,07 | 0,10 | -1,274 | 0,203 | |
| Multip. | Age | <1yo | 1yo | 0,65 | 0,12 | 1,17 | 0,19 | 3,479 | **0,010** | * |
| Strength | | <1yo | 2yo | 0,79 | 0,36 | 1,22 | 0,15 | 5,192 | **0,001** | ** |
| | | 1yo | 2yo | 0,14 | -0,25 | 0,53 | 0,14 | 1,008 | 0,335 | |
| | Sex | Female | Male | 0,14 | -0,18 | 0,45 | 0,14 | 0,952 | 0,361 | |
| | MatR | Low | High | -0,42 | -0,66 | -0,17 | 0,11 | -3,751 | **0,003** | ** |
| | Group | KB | AK | 0,02 | -0,22 | 0,27 | 0,11 | 0,215 | 0,834 | |
| Multip. | Age | <1yo | 1yo | 1,16 | -0,48 | 2,79 | 0,68 | 1,697 | 0,269 | |
| Eigenv. | | <1yo | 2yo | 0,83 | -0,44 | 2,09 | 0,53 | 1,564 | 0,269 | |
| | | 1yo | 2yo | -0,33 | -1,66 | 0,99 | 0,55 | -0,597 | 0,550 | |
| | Sex | Female | Male | 0,11 | -0,91 | 1,13 | 0,52 | 0,209 | 0,834 | |
| | MatR | Low | High | -1,47 | -2,23 | -0,71 | 0,39 | -3,784 | **p<0,001** | *** |
| | Group | KB | AK | 0,66 | -0,13 | 1,44 | 0,40 | 1,636 | 0,102 | |

*<u>Note</u>: (1) *p<0.05; **p<0.01; ***p<0.001; (2) SE = Standard Error, CI = Confidence Interval*

**Table S4. Effects of age, sex, maternal rank (MatR) and social group on multiplex metrics (i.e., multiplex degree, strength and eigenvector) of the *social interactions* network.**

| | Pred. | Level1 | Level2 | Diff. | CI_low | CI_high | SE | t/z | p | |
|---|---|---|---|---|---|---|---|---|---|---|
| Multip. | Age | <1yo | 1yo | -0,07 | -0,53 | 0,40 | 0,19 | -0,335 | 0,849 | |
| Degree | | <1yo | 2yo | 0,13 | -0,26 | 0,51 | 0,16 | 0,799 | 0,849 | |
| | | 1yo | 2yo | 0,19 | -0,13 | 0,52 | 0,14 | 1,427 | 0,461 | |
| | Sex | Female | Male | 0,25 | -0,04 | 0,54 | 0,15 | 1,677 | 0,094 | |
| | MatR | Low | High | -0,23 | -0,44 | -0,02 | 0,11 | -2,112 | **0,035** | * |
| | Group | KB | AK | -0,23 | -0,45 | -0,01 | 0,11 | -2,094 | **0,036** | * |
| Multip. | Age | <1yo | 1yo | 0,08 | -0,11 | 0,27 | 0,07 | 1,178 | 0,527 | |
| Strength | | <1yo | 2yo | 0,10 | -0,05 | 0,26 | 0,05 | 1,912 | 0,247 | |
| | | 1yo | 2yo | 0,03 | -0,11 | 0,16 | 0,05 | 0,509 | 0,621 | |
| | Sex | Female | Male | 0,18 | 0,06 | 0,29 | 0,05 | 3,476 | **0,005** | ** |
| | MatR | Low | High | -0,06 | -0,14 | 0,03 | 0,04 | -1,413 | 0,185 | |
| | Group | KB | AK | 0,02 | -0,07 | 0,11 | 0,04 | 0,440 | 0,669 | |
| Multip. | Age | <1yo | 1yo | 2,65 | 0,82 | 4,47 | 0,76 | 3,472 | **0,002** | ** |
| Eigenv. | | <1yo | 2yo | 1,96 | 0,49 | 3,42 | 0,61 | 3,201 | **0,003** | ** |
| | | 1yo | 2yo | -0,69 | -1,95 | 0,57 | 0,53 | -1,315 | 0,188 | |
| | Sex | Female | Male | 1,55 | 0,41 | 2,70 | 0,59 | 2,652 | **0,008** | ** |

| | | | | | | | | | |
|---|---|---|---|---|---|---|---|---|---|
| MatR | Low | High | 1,42 | 0,51 | 2,33 | 0,46 | 3,058 | **0,002** | ** |
| Group | KB | AK | 0,53 | -0,29 | 1,35 | 0,42 | 1,268 | 0,205 | |

*Note: (1) *p<0.05; **p<0.01; ***p<0.001; (2) SE = Standard Error, CI = Confidence Interval*

#### 3. Network consistency across different ages

##### *3.1. Kin-proximity-Social*

**Table S5. Age groups comparisons of edge overlap between proximity and social networks**

| Compared | Networks | Level1 | Level2 | Diff. | CI_low | CI_high | SE | z | p | |
|---|---|---|---|---|---|---|---|---|---|---|
| Kinship - | Proximity | 1yo | <1yo | -0,07 | 0,02 | -0,12 | -0,02 | -2,82 | **0,005** | ** |
| | | 2yo | <1yo | -0,11 | 0,02 | -0,15 | -0,06 | -4,32 | **p<0,001** | *** |
| | | 2yo | 1yo | -0,04 | 0,02 | -0,08 | 0,01 | -1,54 | 0,123 | |
| Kinship - | Social interactions | 1yo | <1yo | 0,01 | 0,01 | -0,01 | 0,03 | 0,91 | 0,361 | |
| | | 2yo | <1yo | -0,02 | 0,01 | -0,04 | 0,00 | -2,29 | **0,022** | * |
| | | 2yo | 1yo | -0,03 | 0,01 | -0,04 | -0,01 | -3,18 | **0,001** | ** |
| Proximity - | Social interactions | 1yo | <1yo | 0,09 | 0,03 | 0,04 | 0,15 | 3,27 | **0,001** | ** |
| | | 2yo | <1yo | 0,05 | 0,03 | -0,01 | 0,10 | 1,75 | 0,081 | |
| | | 2yo | 1yo | -0,04 | 0,03 | -0,10 | 0,01 | -1,52 | 0,128 | |

*Note: (1) *p<0.05; **p<0.01; ***p<0.001; (2) SE = Standard Error, CI = Confidence Interval*

##### *3.2. Kin-proximity-grooming-groomed*

**Table S6. Age groups comparisons of edge overlap between kinship and proximity and allogrooming**

| Compared | Networks | Level1 | Level2 | Diff. | CI_low | CI_high | SE | z | p | |
|---|---|---|---|---|---|---|---|---|---|---|
| Kinship - | Grooming | 1yo | <1yo | 0,13 | 0,01 | 0,10 | 0,16 | 9,25 | **p<0,001** | *** |
| | | 2yo | <1yo | 0,08 | 0,01 | 0,06 | 0,10 | 6,68 | **p<0,001** | *** |
| | | 2yo | 1yo | -0,05 | 0,02 | -0,08 | -0,02 | -2,89 | **0,004** | ** |
| | Groomed | 1yo | <1yo | 0,04 | 0,02 | 0,01 | 0,08 | 2,70 | 0,007 | ** |
| | | 2yo | <1yo | 0,03 | 0,02 | 0,00 | 0,06 | 1,97 | **0,049** | * |
| | | 2yo | 1yo | -0,01 | 0,02 | -0,05 | 0,02 | -0,77 | 0,444 | |
| Proximity - | Grooming | 1yo | <1yo | 0,34 | 0,04 | 0,27 | 0,41 | 9,70 | **p<0,001** | *** |
| | | 2yo | <1yo | 0,25 | 0,03 | 0,19 | 0,32 | 7,52 | **p<0,001** | *** |
| | | 2yo | 1yo | -0,09 | 0,04 | -0,18 | 0,00 | -2,03 | **0,042** | * |

| | | | | | | | | | | |
|---|---|---|---|---|---|---|---|---|---|---|
| | Groomed | 1yo | <1yo | 0,08 | 0,03 | 0,02 | 0,14 | 2,51 | **0,012** | * |
| | | 2yo | <1yo | 0,01 | 0,03 | -0,05 | 0,07 | 0,23 | 0,816 | |
| | | 2yo | 1yo | -0,07 | 0,03 | -0,14 | -0,01 | -2,26 | 0,024 | |
| Grooming- | Groomed | 1yo | <1yo | 0,37 | 0,06 | 0,26 | 0,48 | 6,41 | **p<0,001** | *** |
| | Gr-Gd | 2yo | <1yo | 0,32 | 0,06 | 0,21 | 0,43 | 5,57 | **p<0,001** | *** |
| | Gr-Gd | 2yo | 1yo | -0,05 | 0,07 | -0,18 | 0,08 | -0,75 | 0,456 | |
